## Supplementary material for "Senescence in yeast is associated with chromosome XII fragments rather than ribosomal DNA circle accumulation": Table S1

| YJH  1530 | WT | *MAT a/MAT ɑ ade2::hisG/ade2::hisG his3/his3 leu2/leu2 lys2/+ met15D::ADE2/+ ura3D0/ura3Δo trp1D63/trp1D63 hoD::SCW11pr-Cre-EBD78-NatMX/hoD::SCW11pr-Cre-EBD78-NatMX loxP-UBC9-loxP-LEU2/lOXp-UBC9-loxP-LEU2 loxP-CDC20-Intron-loxP-HPHMX/loxP-CDC20-Intron-loxP-HPHMX* | (Lindstrom and Gottschling, 2009)-UCC5185 |
| --- | --- | --- | --- |
| YJH  1069 | *spt3Δ* | *MAT a/MAT ɑ ade2::hisG/ade2::hisG his3/his3 leu2/leu2 lys2/+ met15D::ADE2/+ ura3D0/ura3Δo trp1D63/trp1D63 hoD::SCW11pr-Cre-EBD78-NatMX/hoD::SCW11pr-Cre-EBD78-NatMX loxP-UBC9-loxP-LEU2/lOXp-UBC9-loxP-LEU2 loxP-CDC20-Intron-loxP-HPHMX/loxP-CDC20-Intron-loxP-HPHMX spt3::TRP1/spt3::TRP1* | (Hull et al., 2019) |
| YJH  1052 | *rad52Δ* | *MAT a/MAT ɑ ade2::hisG/ade2::hisG his3/his3 leu2/leu2 lys2/+ met15D::ADE2/+ ura3D0/ura3Δo trp1D63/trp1D63 hoD::SCW11pr-Cre-EBD78-NatMX/hoD::SCW11pr-Cre-EBD78-NatMX loxP-UBC9-loxP-LEU2/lOXp-UBC9-loxP-LEU2 loxP-CDC20-Intron-loxP-HPHMX/loxP-CDC20-Intron-loxP-HPHMX rad52::TRP1/rad52::TRP1* | This study |
| YAZ  47 | *sac3Δ* | *MAT a/MAT ɑ ade2::hisG/ade2::hisG his3/his3 leu2/leu2 lys2/+ met15D::ADE2/+ ura3D0/ura3Δo trp1D63/trp1D63 hoD::SCW11pr-Cre-EBD78-NatMX/hoD::SCW11pr-Cre-EBD78-NatMX loxP-UBC9-loxP-LEU2/lOXp-UBC9-loxP-LEU2 loxP-CDC20-Intron-loxP-HPHMX/loxP-CDC20-Intron-loxP-HPHMX sac3::TRP1/sac3::TRP1* | This study |
| YAZ  38 | TRP1 repaired | *MAT a/MAT ɑ ade2::hisG/ade2::hisG his3/his3 leu2/leu2 lys2/+ met15D::ADE2/+ ura3D0/ura3Δo hoD::SCW11pr-Cre-EBD78-NatMX/hoD::SCW11pr-Cre-EBD78-NatMX loxP-UBC9-loxP-LEU2/lOXp-UBC9-loxP-LEU2 loxP-CDC20-Intron-loxP-HPHMX/loxP-CDC20-Intron-loxP-HPHMX* | This study |
| YCJ  261 | *fob1*Δ | *MAT a/MAT ɑ ade2::hisG/ade2::hisG his3/his3 leu2/leu2 lys2/+ met15D::ADE2/+ ura3D0/ura3Δo trp1D63/trp1D63 hoD::SCW11pr-Cre-EBD78-NatMX/hoD::SCW11pr-Cre-EBD78-NatMX loxP-UBC9-loxP-LEU2/lOXp-UBC9-loxP-LEU2 loxP-CDC20-Intron-loxP-HPHMX/loxP-CDC20-Intron-loxP-HPHMX fob1Δ::kanMX.fob1Δ::kanMX* | (Lindstrom and Gottschling, 2009)- UCC526 |
| YDH  57 | *spt3*Δ TOM70-GFP Rpl13a-mCherry | *MAT a/MAT ɑ ade2::hisG/ade2::hisG his3/his3 leu2/leu2 lys2/+ met15D::ADE2/+ ura3D0/ura3Δo trp1D63/trp1D63 hoD::SCW11pr-Cre-EBD78-NatMX/hoD::SCW11pr-Cre-EBD78-NatMX loxP-UBC9-loxP-LEU2/lOXp-UBC9-loxP-LEU2 loxP-CDC20-Intron-loxP-HPHMX/loxP-CDC20-Intron-loxP-HPHMX TOM70-GFP-TRP1/+ RPL13A-mCherry-KanMX6/+ spt3::URA3/spt3::URA3* | This study |
| YDH  52 | *rad52*Δ TOM70-GFP Rpl13a-mCherry | *MAT a/MAT ɑ ade2::hisG/ade2::hisG his3/his3 leu2/leu2 lys2/+ met15D::ADE2/+ ura3D0/ura3Δo trp1D63/trp1D63 hoD::SCW11pr-Cre-EBD78-NatMX/hoD::SCW11pr-Cre-EBD78-NatMX loxP-UBC9-loxP-LEU2/lOXp-UBC9-loxP-LEU2 loxP-CDC20-Intron-loxP-HPHMX/loxP-CDC20-Intron-loxP-HPHMX TOM70-GFP-TRP1/+ RPL13A-mCherry-KanMX6/+ rad52::URA3/rad52::URA3* | This study |
| YDH  18 | TOM70-GFP Rpl13a-mCherry | *MAT a/MAT ɑ ade2::hisG/ade2::hisG his3/his3 leu2/leu2 lys2/+ met15D::ADE2/+ ura3D0/ura3Δo trp1D63/trp1D63 hoD::SCW11pr-Cre-EBD78-NatMX/hoD::SCW11pr-Cre-EBD78-NatMX loxP-UBC9-loxP-LEU2/lOXp-UBC9-loxP-LEU2 loxP-CDC20-Intron-loxP-HPHMX/loxP-CDC20-Intron-loxP-HPHMX TOM70-GFP-TRP1/+ RPL13A-mCherry-KanMX6/+* | Horkai & Houseley 2021 |
| YJH  1521 | *sac3*Δ TOM70-GFP Rpl13a-mCherry | *MAT a/MAT ɑ ade2::hisG/ade2::hisG his3/his3 leu2/leu2 lys2/+ met15D::ADE2/+ ura3D0/ura3Δo trp1D63/trp1D63 hoD::SCW11pr-Cre-EBD78-NatMX/hoD::SCW11pr-Cre-EBD78-NatMX loxP-UBC9-loxP-LEU2/lOXp-UBC9-loxP-LEU2 loxP-CDC20-Intron-loxP-HPHMX/loxP-CDC20-Intron-loxP-HPHMX TOM70-GFP-TRP1/+ RPL13A-mCherry-KanMX6/+ sac3::URA3/sac3::TRP1* | This study |
| YJH  1540 | *sac3*Δ *rad52*Δ TOM70-GFP Rpl13a-mCherry | *MAT a/MAT ɑ ade2::hisG/ade2::hisG his3/his3 leu2/leu2 lys2/+ met15D::ADE2/+ ura3D0/ura3Δo trp1D63/trp1D63 hoD::SCW11pr-Cre-EBD78-NatMX/hoD::SCW11pr-Cre-EBD78-NatMX loxP-UBC9-loxP-LEU2/lOXp-UBC9-loxP-LEU2 loxP-CDC20-Intron-loxP-HPHMX/loxP-CDC20-Intron-loxP-HPHMX TOM70-GFP-TRP1/+ RPL13A-mCherry-KanMX6/+ sac3::URA3/sac3::TRP1 rad52:HIS3/rad52::HIS3* | This study |
| YJH  1526 | *spt3*Δ *rad52*Δ TOM70-GFP Rpl13a-mCherry | *MAT a/MAT ɑ ade2::hisG/ade2::hisG his3/his3 leu2/leu2 lys2/+ met15D::ADE2/+ ura3D0/ura3Δo trp1D63/trp1D63 hoD::SCW11pr-Cre-EBD78-NatMX/hoD::SCW11pr-Cre-EBD78-NatMX loxP-UBC9-loxP-LEU2/lOXp-UBC9-loxP-LEU2 loxP-CDC20-Intron-loxP-HPHMX/loxP-CDC20-Intron-loxP-HPHMX TOM70-GFP-TRP1/+ RPL13A-mCherry-KanMX6/+ spt3::URA3/spt3::TRP1 rad52:HIS3/rad52::HIS3* | This study |

| YJH  1725 | Tom70-GFP ADE2  GST-V-U | *MAT a/a ade2::hisG/+ his3 leu2 lys2 ura3DO trp1D63 hoD::SCW11pr-Cre-EBD78-NatMX/hoD::SCW11pr-Cre-EBD78-His3MX loxP-UBC9-loxP-LEU2 loxP-CDC20-Intron-loxP-HPHMX Tom70-GFP-[no I-SceI]-KanMX6/+ chrXII:491k-pGST-chrVfrag-ISceIsite-chrVfrag-U* | This study |
| --- | --- | --- | --- |
| YJH  1726 | Tom70-GFP ADE2 XII<>V translocation | *MAT a/a ade2::hisG/+ his3 leu2 lys2 ura3DO trp1D63 hoD::SCW11pr-Cre-EBD78-NatMX/hoD::SCW11pr-Cre-EBD78-His3MX loxP-UBC9-loxP-LEU2 loxP-CDC20-Intron-loxP-HPHMX Tom70-GFP-[no I-SceI]-KanMX6/+ chrXII:1-491k-Pgal-SceI-TRP1--chrV:565k-end, chrV:1-565k--URA3-chrXII:491k-end* | This study |
| YDH  17 | Tom70-GFP VPH1-mCherry | *MAT a/MAT ɑ ade2::hisG his3 leu2 met15D/MET15 lys2/LYS2 ura3DO trp1D63 hoD::SCW11pr-Cre-EBD78-NatMX loxP-UBC9-loxP-LEU2 loxP-CDC20-Intron-loxP-HPHMX Tom70-GFP-TRP1 VPH1-mCherry-Kan* | This study |
| YDH  63 | *rad52*Δ Tom70-GFP VPH1-mCherry | *MAT a/MAT ɑ ade2::hisG his3 leu2 met15D/MET15 lys2/LYS2 ura3DO trp1D63 hoD::SCW11pr-Cre-EBD78-NatMX loxP-UBC9-loxP-LEU2 loxP-CDC20-Intron-loxP-HPHMX Tom70-GFP-TRP1 VPH1-mCherry-Kan rad52::URA3* | This study |
| YDH  66 | *spt3*Δ Tom70-GFP VPH1-mCherry | *MAT a/MAT ɑ ade2::hisG his3 leu2 met15D/MET15 lys2/LYS2 ura3DO trp1D63 hoD::SCW11pr-Cre-EBD78-NatMX loxP-UBC9-loxP-LEU2 loxP-CDC20-Intron-loxP-HPHMX Tom70-GFP-TRP1 VPH1-mCherry-Kan spt3::URA3* | This study |
| YDH  142 | Rpa190-GFP | *MAT a/MAT ɑ ade2::hisG his3 leu2 lys2/+ met15D::ADE2/+ ura3D0/ura3Δo hoD::SCW11pr-Cre-EBD78-NatMX loxP-UBC9-loxP-LEU2 loxP-CDC20-Intron-loxP-HPHM +/RPA190-GFP-HIS3* | This study |
| YJH  1517 | *spt3*Δ  Rpa190-GFP | *MAT a/MAT ɑ ade2::hisG his3 leu2 met15D::ADE2 ura3D0 hoD::SCW11pr-Cre-EBD78-NatMX loxP-UBC9-loxP-LEU2 loxP-CDC20-Intron-loxP-HPHMX RPA190-GFP-HIS3/+ spt3::KAN/spt3::kan* | This study |
| YJH  1518 | *rad52*Δ  Rpa190-GFP | *MAT a/MAT ɑ ade2::hisG his3 leu2 met15D::ADE2 ura3D0 hoD::SCW11pr-Cre-EBD78-NatMX loxP-UBC9-loxP-LEU2 loxP-CDC20-Intron-loxP-HPHMX RPA190-GFP-HIS3/+ rad52::KanMX6/rad52::TRP1* | This study |
| YJH  1709 | *mus81*Δ  TOM70-GFP Rpl13a-mCherry | *MAT a/MAT ɑ ade2::hisG his3 leu2 met15D::ADE2/+ lys2/+ ura3D0 trp1D63 hoD::SCW11pr-Cre-EBD78-NatMX loxP-UBC9-loxP-LEU2 loxP-CDC20-Intron-loxP-HPHMX Tom70-GFP-TRP1/+ RPL13A-mCherry-Kan/+ mus81::URA3/mus81::TRP1* | This study |
| YHM  30 | *hst3*Δ *hst4*Δ  TOM70-GFP Rpl13a-mCherry | *MAT a/MAT ɑ ade2::hisG his3 leu2 met15D::ADE2/+ lys2/+ ura3D0 trp1D63 hoD::SCW11pr-Cre-EBD78-NatMX loxP-UBC9-loxP-LEU2 loxP-CDC20-Intron-loxP-HPHMX Tom70-GFP-TRP1/+ RPL13A-mCherry-Kan/+ hst3::URA3 hst4::HIS* | This study |
