## Supplementary material for "Senescence in yeast is associated with chromosome XII fragments rather than ribosomal DNA circle accumulation": Table S2

| oJH1352 | SPT3 UP45 | GTGCATACAAGGTACCGTGCCAAAATTCAAGAGATTAGGGCAGGA CGGATCCCCGGGTTAATTAAG |
| --- | --- | --- |
| oJH1353 | SPT3 DN45 | CCAGAAGGAAACCCATGCACCTCCATGATGAAATTATACAAAAAT GAATTCGAGCTCGTTTAAAC |
| oJH1354 | SPT3 C | ACCCCATGATGATGTGATTG |
| oJH1355 | SPT3 D | GCCCCATAGTATCTCAACATCA |
| oJH1327 | RAD52 UP45 2 | TTGCCAAGAACTGCTGAAGGTTCTGGTGGCTTTGGTGTGTTGTTGGAATTCGAGCTCGTTTAAAC |
| oJH1328 | RAD52 DN45 2 | TAAATAATGATGCAAATTTTTTATTTGTTTCGGCCAGGAAGCGTTCGGATCCCCGGGTTAATTAAG |
| oJH1239 | RAD52 C2 | TAGGCACACCGTTGATCAGA |
| oJH1240 | RAD52 D2 | CACATGGAGGAAAGAAAAACTAGA |
| oAW88 | SAC3 UP45 | TGTTGGTACTCATTTCAGGTAATACTCTTGGAAGGTATCCCTTAACGGATCCCCGGGTTAATTAAG |
| oAW89 | SAC3 DN45 | TATAGAAAAAATGCACATTTCTTTTGTTTATATATTACAAATGCTGAATTCGAGCTCGTTTAAAC |
| oAW90 | SAC3 C | CCGTATTACCCGTTAGCACA |
| oAW91 | SAC3 D | CGCCGACAGTAATAAGAAATG |
| oAZ110 | TRP1 UP | CATGGAGGGCGTTATTACG |
| oAZ111 | TRP1 DN | CCGCTAATAGGAAGTATGAATACC |
| oAZ112 | TRP1 C | GAGACCGAGTTAGGGACAGTTAG |
| oAZ113 | TRP1 D | AGCGAAAAGACGATAAATACAAG |
| oJH1520 | TOM70 DN45 1 | TAGTTTTTGTCTTCTCCTAAAAGTTTTTAAGTTTATGTTTACTGT GAATTCGAGCTCGTTTAAAC |
| oJH1521 | TOM70 UP45 1 | AAGATTCAAGAAACTTTAGCTAAATTACGCGAACAGGGTTTAATG CGG ATC CCC GGG TTA ATT AAC |
| oJH1522 | TOM70 C | AAGTTGGATGGTATTTATGTTGGA |
| oJH1523 | TOM70 D | GAACATGCATACGTGTATATTGAAAC |
| oDH11 | RPL13A-mCherry UP45 | AAGAGAGCTAGAGAAAAGGCTGAAGCTGAAGCTGAAAAGAAGAAATGCATGCTTATGGTGAGCAA |
| oDH12 | RPL13A-mCherry DN45 | ATACAAAAATTGTGGATGAAAAATTCTTTGATGAAGTTTTTAGATAAGTTATACTAGTTCGTCGACTGGAT |
| oDH8 | RPL13A C | CAGAACCTTGAGATTAGCCAGAT |
| oDH9 | RPL13A D | ATCTTTCGCATCTCTTCTATGC |
| oJH1819 | pFA6a I-SceI fix F | ATAGATCTGTTTAGCTTGCCTCG |
| oJH1820 | pFA6a I-SceI fix R | GCGGATCTGCCGGTAGAG |
| oJH1808 | ChrXII insert UP45 1 | gtgtttatcagatcagtgtgatgtgttcagctaaatggaaagcta CCGCGCGTTGGCCGATTCAT |
| oJH1809 | ChrXII insert DN45 1 | tactgtgagaccttttcttgcaatatctcgatcagatagcatctg TTCGTACGCTGCAGGTCGAC |
| oJH1798 | Chr V-XII inversion F1 | Gtcacattga GAGCTCCCGCTCTTTTTCAAAC |
| oJH1799 | Chr V-XII inversion R1 | Gtcacattga ggatc CGATACAGTGTTCCAGAATTGTCA |
| oJH1800 | Chr V-XII inversion F2 | Gtcacattga ggatcc TAGGGATAACAGGG TAATCCCAAATAATGTATGTAGAATAGA |
| oJH1801 | Chr V-Xii inversion R2 | Gtcacattga CCGCGG AGGTATGTCATCGTAATCAAGT |
| oDH1 | VPH1 UP45 1 | GACATGGAAGTCGCTGTTGCTAGTGCAAGCTCT TCC GCT TCA AGC TGC ATG CTT ATG GTG AGC AA |
| oDH2 | VPH1 DN45 1 | AATGAAGTACTTAAATGTTTCGCTTTTTTTAAAAGTCCTCAAAAT AAGTTATACTAGTTCGTCGACTGGAT |
| oAZ114 | RPA190 tag UP45 | GGTACGGGTTCATTTGATGTGTTAGCAAAGGTTCCAAATGCGGCT CGGATCCCCGGGTTAATTAAC |
| oAZ115 | RPA190 tag DN45 | AAACTAATATTAAATCGTAATAATTATGGGACCTTTTGCCTGCTT GAATTCGAGCTCGTTTAAAC |
| oRH1 | Mus81 F1 | AAGTTTCAAAGGATTGATACGAACACACATTCCTAGCATGAAAGCCGGATCCCCGGGTTAATTAAG |
| oRH2 | Mus81 R1 | TTTTTTCTTTATAAAACCTTGCAGGGATGACTATATTTCAAATTGGAATTCGAGCTCGTTTAAAC |
| oCJ124 | Hst3 up | AAGGGATTAATTTACATACAACTAGATCCATCTTTCTCAAAAATGCGGATCCCCGGGTTAATTAAG |
| oCJ125 | Hst3 dn | TGCCTCGATTATTTATCGTTAACTCAATTTTAATAGTTAAGTTTAGAATTCGAGCTCGTTTAAAC |
| oCJ11 | HST4 dn 45 r1 cj | TGCATTAATTTTATCTCCAACCTTTTTTGGTAGGACAAATACTTTGAATTCGAGCTCGTTTAAAC |
| oCJ12 | HST4 up45 f 1 cj | CAACTGTATTTTAAAACTGTAAATAATACTAAGCAGAACCACAAACGGATCCCCGGGTTAATTAAG |
