## Supplementary material for "Senescence in yeast is associated with chromosome XII fragments rather than ribosomal DNA circle accumulation": Table S3

| pJH9 | pFA6a TRP1 | (Longtine et al., 1998) |
| --- | --- | --- |
| pJH20 | pFA6a GFP HIS3MX6 | (Longtine et al., 1998) |
| pJH369 | pAW8-mCherry | (Watson et al., 2008) |
| pJH388 | pFA6a-GFP-KanMX6 no I-SceI | I-SceI site removed by site directed mutagenesis |
| pJH381 | pGSTU | delitto perfetto plasmid with Kan swapped out for TRP1 |
| pJH387 | pGST-V-U | For reciprocal translocations with Chr V sub-telomere. Has region of V:564025-566025 inserted between URA3 and TRP1 markers with SceI site in the middle. Note plasmid yield was weirdly low |
| pJH73 | pFA6a URA3 | (Houseley and Tollervey, 2011) |
