## Supplementary figures and images for "Senescence in yeast is associated with chromosome XII fragments rather than ribosomal DNA circle accumulation"

### File S4

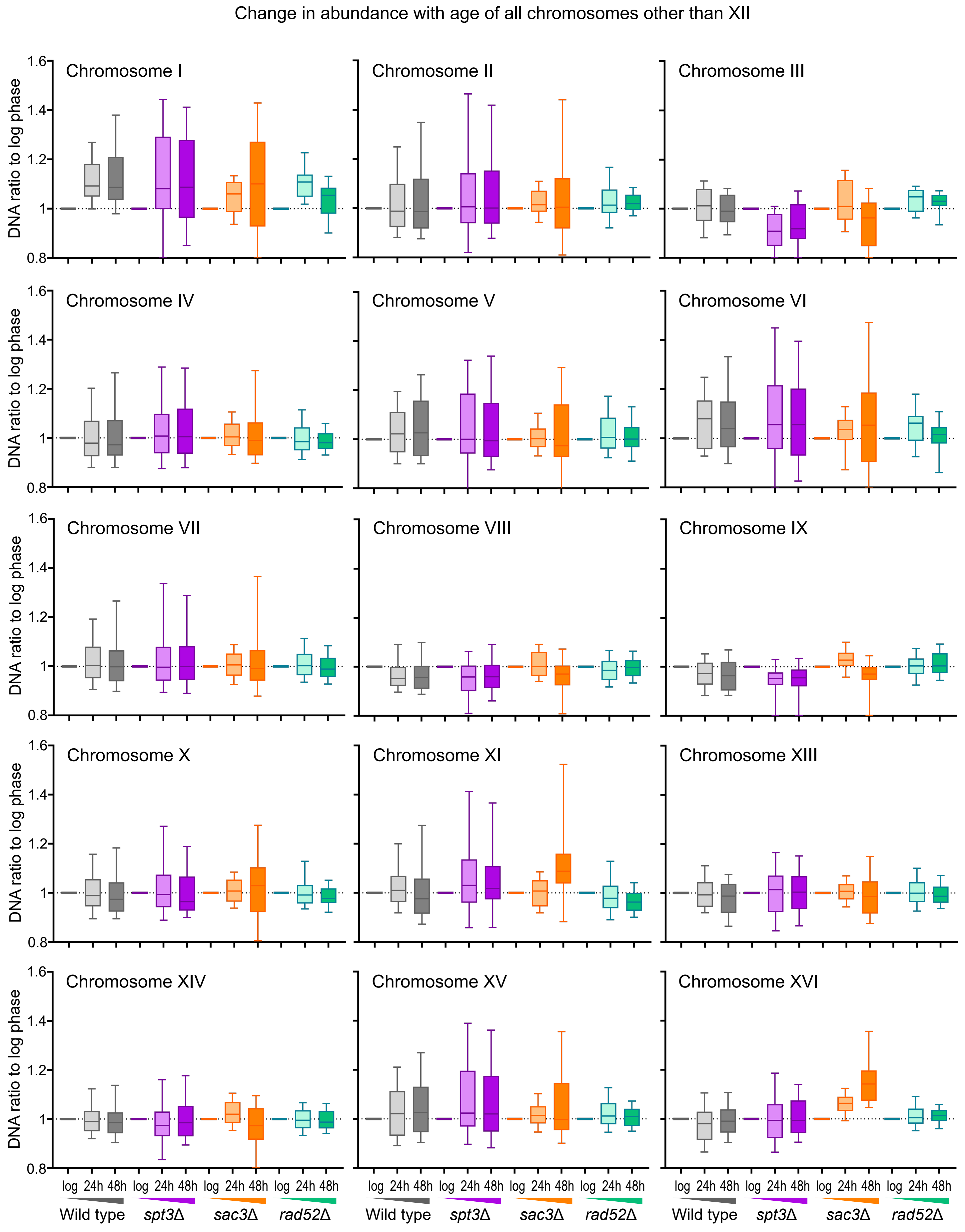
